# Absence of CD47 in the tumor microenvironment modulates tumor metabolism and immunosuppressive signatures limiting breast cancer progression

**DOI:** 10.1101/2023.07.12.548766

**Authors:** Elizabeth R. Stirling, Yu-Ting Tsai, Steven M. Bronson, Adam Wilson, Brian Westwood, Alexandra Thomas, Pierre L. Triozzi, Cristina M. Furdui, Glenn J. Lesser, Katherine L. Cook, David R. Soto-Pantoja

## Abstract

The majority of breast cancers are generally considered immune-deprived tumors. This lack of immunogenicity severely hinders effectiveness of current immunotherapy approaches limiting therapeutic options to control disease. Therefore, we need new biomarkers to determine and enhance immune responses to improve the outcome of cancer patients experiencing invasive disease. Our data in matched human patient biopsies show that CD47 expression increases from primary to metastatic tumors. CD47 is an integral membrane protein that impairs antitumor immunosurveillance and influences normal tissue metabolism. However, whether CD47 plays a role in regulating tumor bioenergetics is unknown. A carcinogen-induced mouse mammary carcinogenesis model demonstrates that the absence of CD47 reduces tumor burden, which is associated with a distinct metabolic signature compared to WT tumors. Depletion of several lipid metabolites was observed in the absence of CD47, and metabolic dependency experiments suggest that anti-sense blockade of CD47 limits reliance on fatty acid oxidation as a fuel supporting cellular respiration on cancer cells. Our global metabolomics analysis also implicated the absence of CD47 in downregulation of immunosuppressive metabolites of the tryptophan and prostaglandin pathways. Spatial proteomic analysis revealed increased immune infiltrate and substantial reduction in immunosuppressive immune checkpoint proteins in the absence of CD47 with the highest reduction in intra-tumoral PD-L1 expression. Since anti-PD-L1 therapy is used in the current strategy to treat triple-negative breast cancer (TNBC), we targeted CD47 in an EMT-6 syngeneic TNBC model. The *in vivo* knockdown of CD47 sensitized tumors to anti-PD-L1 therapy to decrease tumor burden and increase intratumoral cytotoxic T cells. Therefore, targeting CD47 may be a suitable immunotherapeutic option to limit immunosuppression and enhance the efficacy of immune checkpoint blockade.

## Introduction

Invasive breast cancer is the second leading cause of cancer death among women in the United States, with over 200,000 expected cases annually. Disease progression is associated with the development of therapeutic resistance, allowing re-growth of the tumor and metastatic spread of cancer cells. It is well recognized that metabolism plays a critical role in fueling cancer cells energy capacity to proliferate and spread. Aside from supporting cell growth, specific metabolites affect immune signaling in the tumor microenvironment resulting in immunosuppression and lack of response to current immunotherapy approaches. Therefore, strategies that interfere with cancer cell metabolite consumption and limit immunosuppression could be attractive to overcome breast cancer invasiveness and enhance antitumor immunity.

CD47 is a ubiquitously expressed type one integral membrane protein that is often overexpressed in both hematological and solid tumor cancers, impairing both antitumor innate and adaptive immune surveillance ^4–6^. When CD47 on cancer cells interacts with the innate immune cell counterreceptor signal regulatory protein alpha (SIRPα), antiphagocytic “don’t eat me” signals arise, allowing cancer cells to bypass immunosurveillance ^7^. Additionally, the ligation of CD47 on a T cell by the matricellular glycoprotein thrombospondin-1 (TSP1) can impair T cell activation, differentiation, and survival ^8–11^. Therefore, CD47 plays an essential role in cancer cell survival as its ligation can impair both innate and adaptive antitumor immune response and allow the cell to evade immune detection.

CD47 has become an attractive cancer immunotherapeutic target as its ligation, and subsequent signaling impairs both innate and adaptive immune cell antitumor response. CD47 blockade decreases primary tumor size and metastasis in several pre-clinical models due to enhanced antitumor immune cell function ^12–14^. Aside from its role in regulating immune cell effector function, CD47 blockade preserves tissue viability during irradiation due to the upregulation of several metabolic pathways ^15, 16^. While the evidence of CD47 regulation of immune responses and metabolism is strong, very little information is known on whether these two aspects are regulated by CD47 in the tumor microenvironment.

This new report examines how the absence of CD47 modulates metabolism and inflammatory signaling in the tumor microenvironment, which reduces breast cancer invasiveness. The breast tumor and mammary gland tissue from this study underwent metabolomic analysis to show that *cd47*-/- breast tumor tissue differentially regulates metabolites associated with proliferation, fatty acid metabolism, inflammation, and immunosuppression compared to WT breast tissue. Collectively, changes in these pathways illustrate the link between metabolic reprogramming in tumor tissue and how CD47 might influence neoplastic transformation. Spatial proteomics in our study showed that CD47 expression mediates upregulation of immunosuppressive molecules in the tumor microenvironment, with PD-L1 expression being the most elevated. Anti-PD-L1 therapy is considered part of the therapeutic regimen to treat Triple-negative breast cancer^3^ (TNBC). Furthermore, It is reported that patients with TNBC are more likely to experience a visceral metastasis within five years of diagnosis than those with other types of cancer ^1^. Therefore, we executed a TNBC model to test whether CD47 blockade would enhance anti-PD-L1 therapy. Our data show that tumor ablation is enhanced by targeting these two receptors. Therefore, blockade of CD47 decreases intratumoral immunosuppressive molecules and enhances immunogenicity allowing tumors to be sensitized to immune checkpoint blockade to reduce TNBC tumor burden.

## Methods

### Carcinogen-induced triple-negative breast cancer mouse model

A carcinogen-induced mouse model was performed to determine the impact of CD47 receptor expression on tumor burden. Female WT and *cd47*-/- (B6.129S7-*Cd47*^tm1Fpl^/J) C57Bl/6 mice were purchased from Jackson Laboratory (n=14-15/group). At 6 weeks of age, mice received 15 mg medroxyprogesterone acetate (MPA) subcutaneously into the mammary gland to stimulate cell proliferation and make the area susceptible to carcinogenesis ^21^. Through week 7-10, mice received oral gavage treatments of 1 mg dimethylbenzathracene (DMBA) suspended in peanut oil once a week ^21^. DMBA is a carcinogen that will promote the growth of breast tumors within the MPA-treated mammary glands. Tumors were palpated and measured twice a week to determine tumor area (LW^2^/2). In contrast, tumor wet weight, incidence, and multiplicity were determined 15 weeks post-DMBA treatment (week 25) when the mice were euthanized.

### Metabolomic Profiling Analysis

Tumors and mammary glands were harvested, flash-frozen, and stored at −80°C until processed (n=9/group). Metabolite analysis was then performed by Metabolon as previously described ^15, 16^. The Metabolon Laboratory Information Management System (LIMS) assigned each sample a unique identifier associated with the original source identifier to track sample handling, tasks, and results.

The automated MicroLab STAR® system from Hamilton Company was used to prepare samples. Before the extraction, recovery standards were added for QC purposes. Small molecules bound to protein or trapped in the precipitated protein matrix were dissociated to remove proteins. To recover chemically diverse metabolites, proteins were precipitated under the aqueous methanol extraction process. This resulted in extract that was divided into five fractions: two for analysis by two separate reverse phase (RP)/UPLC-MS/MS methods with positive ion mode electrospray ionization (ESI), one for analysis by RP/UPLC-MS/MS with negative ion mode ESI, one for analysis by HILIC/UPLC-MS/MS with negative ion mode ESI, and one sample was reserved for backup. Organic solvent was removed by placing samples on a TurboVap® (Zymark). The sample extracts were stored overnight under nitrogen before preparation for analysis.

### Ultrahigh Performance Liquid Chromatography-Tandem Mass Spectroscopy (UPLC-MS/MS)

A Waters ACQUITY UPLC and a Thermo Scientific Q-Exactive mass spectrometer interfaced with heated ESI and Orbitrap mass analyzer were used for all methods. Each sample extract was dried and reconstituted in solvents compatible with each of the four methods. A series of standards at fixed concentrations were used to ensure injection and chromatographic consistency for each reconstituted solvents. Two aliquots were analyzed using acidic positive ion conditions, with one chromatographically optimized for more hydrophilic compounds and the other for hydrophobic conditions. Both aliquots were eluted from a C18 column using water and methanol with 0.05% perfluoropentanoic acid (PFPA) and 0.1% formic acid (FA) for optimizing hydrophilic compounds and acetonitrile, water, 0.05% PFPA and 0.01% FA with operation at an overall higher organic content for optimizing hydrophobic compounds. The third aliquot was analyzed using basic negative ion optimized conditions using a separate dedicated C18 column that eluted extracts from the column using methanol and water with 6.5mM Ammonium Bicarbonate at pH 8. The fourth aliquot was analyzed through negative ionization following elution from a HILIC column using a gradient consisting of water and acetonitrile with 10mM Ammonium Formate at pH 10.8. Alternation between MS and data-dependent MSn scans using dynamic exclusion was performed for MS analysis.

### Bioinformatics

Data analysts used four informatics systems: LIMS, data extraction, and peak-identification software, data processing tools for QC and compound identification, and a collection of information interpretation and visualization tools. LAN backbone and a database server running Oracle 10.2.0.1 Enterprise Edition were used as hardware and software foundations for informatics components.

### Immunohistochemistry

Breast tumors from the carcinogen-induced and EMT-6 TNBC models were harvested, fixed in 4% paraformaldehyde and embedded in paraffin so 5 μM tumor tissue sections could be created with a microtome. The sections were then deparaffinized with xylene three times for 5 minutes and rehydrated with 100% ethanol two times for 5 minutes, 95% ethanol for 3 minutes and 70% ethanol for 3 minutes. Antigen retrieval was performed with 1x citrate buffer. We examined cell proliferation in the carcinogen-induced tumors by examining Ki67 expression (Cell Signaling, Cat #12202S), CD47 expression within a matched human primary and metastatic tumor tissue array (Anti-human CD47 purified (Clone B6H12), eBioscience, Ref #1404782, Lot #4281568) and granzyme B within EMT-6 tumors (Thermo Scientific, Product # PA1-37799, Lot # PA1813162). According to the manufacturer’s protocol, the sections underwent DAB staining (Dako Envision Dual Linked System HRP, Catalog #K4065) and were stained with Ki67, CD47, or granzyme b antibody at a 1:100 dilution. All slides were mounted with Cytoseal XYL (Thermo Fisher, Ref #8312-4). Additionally, EMT-6 tumor sections were stained with anti-mouse CD3 (APC (red), Clone #172A, Catalog #100235), anti-mouse CD8 (Alexa Fluor 488 (green), Clone #53-6.7, Catalog #100726) at a 1:100 dilution to examine cytotoxic T cells with DAPI (blue) denoting nuclei (Thermo Fisher, Ref # 62248, Lot # VG3036772). Cytotoxic T cells were determined and counted based on co-localization of CD3+ and CD8+ cells (yellow). All slides were mounted with ProLongTM Gold Antifade Mountant (ThermoFisher, Catalog #P36930). The Olympus BX43 microscope obtained images while PerkinElmer Mantra and inform software were used for analysis.

### Lipidomic analysis

EMT-6 tumors were analyzed on a Q Exactive HF hybrid quadrupole-Orbitrap mass spectrometer (Thermo Scientific, Waltham, MA, USA) with heated electrospray ionization (HESI) source and a Vanquish UHPLC system (Thermo Scientific, Waltham, MA, USA). Lipids were separated on an Accucore C30 column (2.6µm, 3mm x 150mm, Thermo Scientific, Waltham, MA, USA) using a linear gradient with 60:40 acetonitrile/water (mobile phase A) and 90:10 isopropyl alcohol/acetonitrile (mobile phase B) both of which contain 0.1% formic acid and 10 mM ammonium formate. MS spectra were acquired by data-dependent scans in negative ion mode where MS1 scan identified top ten most abundant precursor ions followed by MS2 where product ions were generated from selected precursor ions. High-energy collisional dissociation (HCD) were utilized for fragmentation with stepped collision energy of 25/30 eV and 30/50/100 eV in each positive and negative polarity mode. Dynamic exclusion was enabled during data-dependent scans. The LC-MS/MS data containing high resolution MS and data dependent MS2 were searched using the LipidSearch v4.2 (Thermo Scientific, Waltham, MA) and search parameters were as follows: precursor mass tolerance, 5 ppm; product mass tolerance, 5 ppm; and species selection. Relative quantification was performed using peak areas normalized to the total ion current (TIC).

### Bioenergetic Analysis

We performed a Seahorse Mito Fuel Flex Test to determine the metabolic pathway dependency for mitochondrial oxidation of EMT-6 mouse breast cancer cells based on CD47 expression. 5,000 EMT-6 cells were seeded into a Seahorse 96 well plate. After the cells adhered for 24 hours at 37°C with 5% CO_2_, EMT-6 cells were treated with 1.5 μL/mL endoporter and 10 μM control (CTRLM) or CD47 morpholino (CD47M). CD47M is an oligonucleotide antisense morpholino that will decrease protein expression of CD47 on the surface of EMT-6 cells. Cells were incubated for 48 hours at 37°C with 5% CO_2_. To determine which metabolic pathway the cells are dependent on for mitochondrial oxidation, sequential injections of 2 μM UK5099, 3 μM BPTES, and/or 4 μM etomoxir occurred. UK5099 obstructs the mitochondrial pyruvate carrier to inhibit glucose oxidation. BPTES prevents glutamine oxidation by inhibiting glutaminase from converting glutamine to glutamate. Etomoxir inhibits carnitine palmitoyl transferase 1A (CPT1A) from transporting long-chain fatty acids into the mitochondria for fatty acid oxidation. Data based on mitochondrial oxidation was expressed as oxygen consumption rate (OCR). These OCR values were used to determine percent dependency by performing the following calculation:

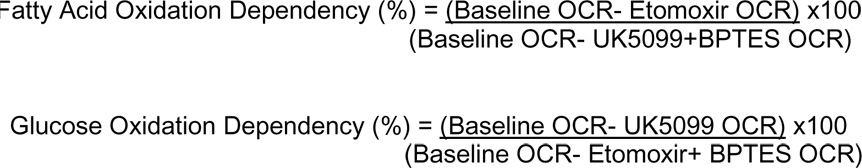

Cells were counted using Brightfield microscopy on the BioTek Cytation 1 Cell Imaging Multi-Mode Reader so the data could be normalized and analyzed with the Wave software.

To determine the glycolytic flux of EMT-6 breast cancer cells, the Seahorse XF96 analyzer was used. 10,000 EMT-6 were seeded into a Seahorse 96 well plate and adhered overnight at 37°C with 5% CO_2_. CD47 was then targeted with 10 μM CD47M and incubated for 48 hours at 37°C with 5% CO_2_. Glycolytic flux was determined by sequential injections of 10 mM glucose, 1 μM oligomycin and 50 mM 2 Deoxy-D-glucose (2-DG). The measurements from this assay are expressed as extracellular acidification rate (ECAR) and analyzed with the Wave software.

### Spatial Proteomic Analysis

To examine protein expression within carcinogen-induced tumors, spatial proteomic analysis was performed using the NanoString *GeoMx*® Digital Spatial Profiler (DSP). Tumors were harvested, fixed in 4% paraformaldehyde, embedded in paraffin, and sectioned into 5 μM tissue sections. Tumor sections were stained with fluorescent antibodies for lymphocytes (CD45+, red), T cells (CD3+, yellow), cancer cells (PanCK, green), and nuclei (DAPI, blue). These antibodies are covalently bound photocleavable DNA indexing oligonucleotides. Regions of interest were exposed to UV light released oligonucleotides and collected for quantification. Data were normalized based on the region of interest.

### Orthotopic triple-negative breast cancer mouse model

To examine if CD47 targeted therapy sensitizes tumors to anti-PD-L1 treatment, an orthotopic TNBC model was performed. Orthotopic injections of 5×10^5^ EMT-6 cells occurred into the right 4/5 mammary fat pad of balb/c mice at 6-8 weeks of age (Jackson Laboratory). When tumors reached 100 mm^3^, mice received alternating day intraperitoneal treatments of 10 µM CD47M (Gene Tools, Philomath, OR) and/or 200 μg anti-PD-L1 therapy (anti-mouse PD-L1 (B7-H1), Bio X Cell, Cat# BE0101, Lot# 720619F1) over 6 days (n=8/group). CD47M treatment will decrease CD47 expression on the EMT-6 tumors. The exact schedule was followed for the controls of 10 µM CTRLM (Gene Tools, Philomath, OR) and/or 200 μg IgG (Mouse IgG2b Isotype Control, Bio X Cell, Cat #BE0088, Lot# 64541901) (n=4-7/group). Tumors were measured every three days with calipers to determine tumor volume. Mice were euthanized when tumors reached 1500 mm^3^ or 21 days when the study ended.

### Statistics

Statistical analysis was performed using significance tests and classification analysis. Significance tests include Welch’s two-sample T-test and/or matched paired T-test for pairwise comparison and one-way, two-way, and/or two-way repeated-measures analysis of variance (ANOVA) for other statistical analysis. The tables display statistically significant metabolites that are downregulated (p<0.05, green and 0.1>p>0.05, light green) and upregulated (p<0.05, red and 0.1>p>0.05, pink). Statistical analysis was performed in ArrayStudio, R (http://cran.r-project.org/), JMP, and GraphPad Prism.

## Results

### Lack of CD47 receptor decreases invasive breast cancer tumor burden

Human primary and matched metastatic tumors were stained to examine CD47 expression **(Figure 1A)**. CD47 was expressed in tumors with increased expression as disease progressed from primary to metastatic tumor, making it an intriguing therapeutic target **(Figure 1B)**. CD47 blockade decreases tumor burden in various cancer models ^12–14^. Therefore, we examined how CD47 receptor expression impacted tumor burden within a carcinogen-induced mouse model. Treatments of MPA and DMBA primed and promoted the growth of tumors **(Figure 1C).** Over the 15 weeks post-DMBA treatment, a decrease in tumor incidence was detected in *cd47*-/- mice compared to WT mice **(Figure 1D)**. When all tumors were accounted for, a reduction in tumor multiplicity was observed in *cd47*-/- mice compared to WT mice **(Figure 1E)**. While differences in tumor multiplicity and latency until tumor formation existed between groups, the morphologic characteristics of the tumors were similar. The tumors were complex, with both groups exhibiting benign and malignant histologic features and a wide range of patterns. Low-grade lesions were characterized by well-organized glandular patterns, and higher-grade tumors were pleomorphic and had less organized glands with poorly demarcated edges. Tumor types included alveolar adenoma **(Supplementary Figure 1A)**; squamous cell carcinomas with multifocal brightly eosinophilic keratin lamination (keratin pearls) **(Supplementary Figure 1B)**; complex carcinoma composed of epithelial cells surrounded by streams of spindle cells **(Supplementary Figure 1C)**; and solid mammary gland carcinoma composed of solid cords of cells with little to no gland formation **(Supplementary Figure 1D)**. Still, the tumors within the *cd47*-/- mice decreased both tumor weight and area compared to WT mice **(Figure 1F-G)**, thus suggesting that the absence of CD47 delays tumor formation and reduces tumor burden in the DMBA model of mammary carcinogenesis.

**Figure 1:**
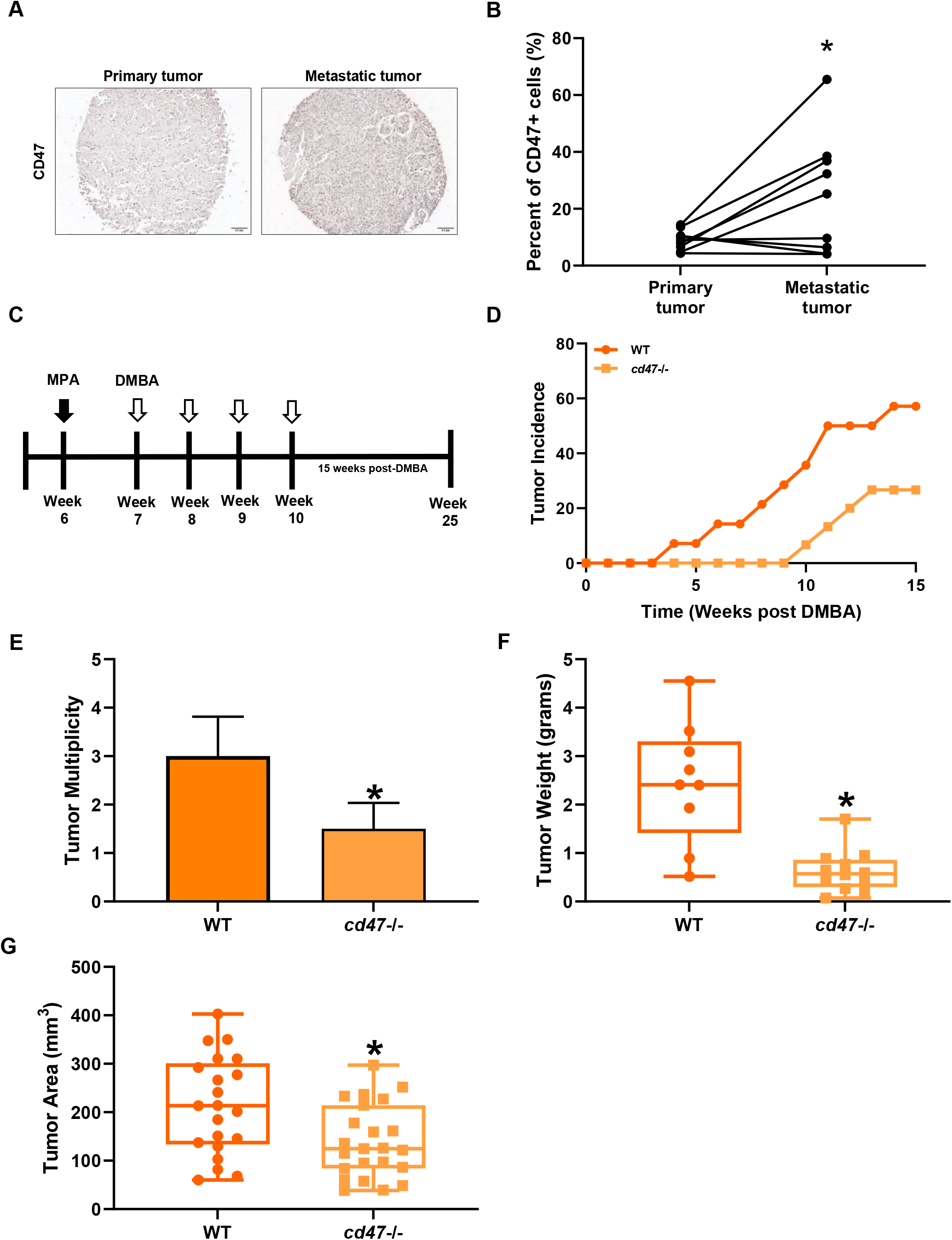
Lack of CD47 receptor decreases tumor burden in a carcinogen-induced triple-negative breast cancer mouse model. **(A)** Representative images of CD47 expression within human matched primary and metastatic tumors. **(B)** Both primary and matched metastatic tumors express CD47, increasing expression as the disease progresses. (*p<0.05, n=8/group) **(C)** Schematic of carcinogen-induced mouse model treatment regimen. At week 6 of age, female WT and *cd47*-/- C57Bl/6 mice were injected subcutaneously with MPA into the mammary fat pad. During weeks 7-10, oral gavages of DMBA were administered once a week. Mice were euthanized on week 25. *cd47*-/- mice decreased **(D)** tumor incidence and **(E)** multiplicity compared to WT mice. Tumors from *cd47*-/- mice decreased **(F)** tumor weight and **(G)** tumor area compared to WT mice. (*p<0.05, n=14-15/group)

### Distinct biochemical signatures of breast tumor and mammary gland tissue are dependent on the expression of CD47

Over 800 compounds were identified within the breast tumor and mammary gland tissue from WT and *cd47*-/- mice. To identify biochemicals that significantly differ between tissue types and genotypes, the following was performed: log transformations, imputation of missing values, if any, with the minimum observed value for each compound, and ANOVA contrasts. A principal component analysis was developed based on the global metabolic profiles of the tissue. The principal component analysis displayed separation between both tissue types and their respective genotype **(Figure 2A)**. Additionally, 162 and 154 metabolites of breast tumors and mammary glands, respectively, were significantly different between genotypes (p<0.05, **Figure 2B, 2D)** with further analysis alternatively displaying metabolites achieving statistical significance of p≤0.01 **(Figure 2F-G)**. Of these statistically significant metabolites, unique biochemical signatures were observed between WT and *cd47*-/- breast tumors and mammary glands regarding 7-8 different metabolic pathways **(Figure 2C, 2E)**.

**Figure 2.**
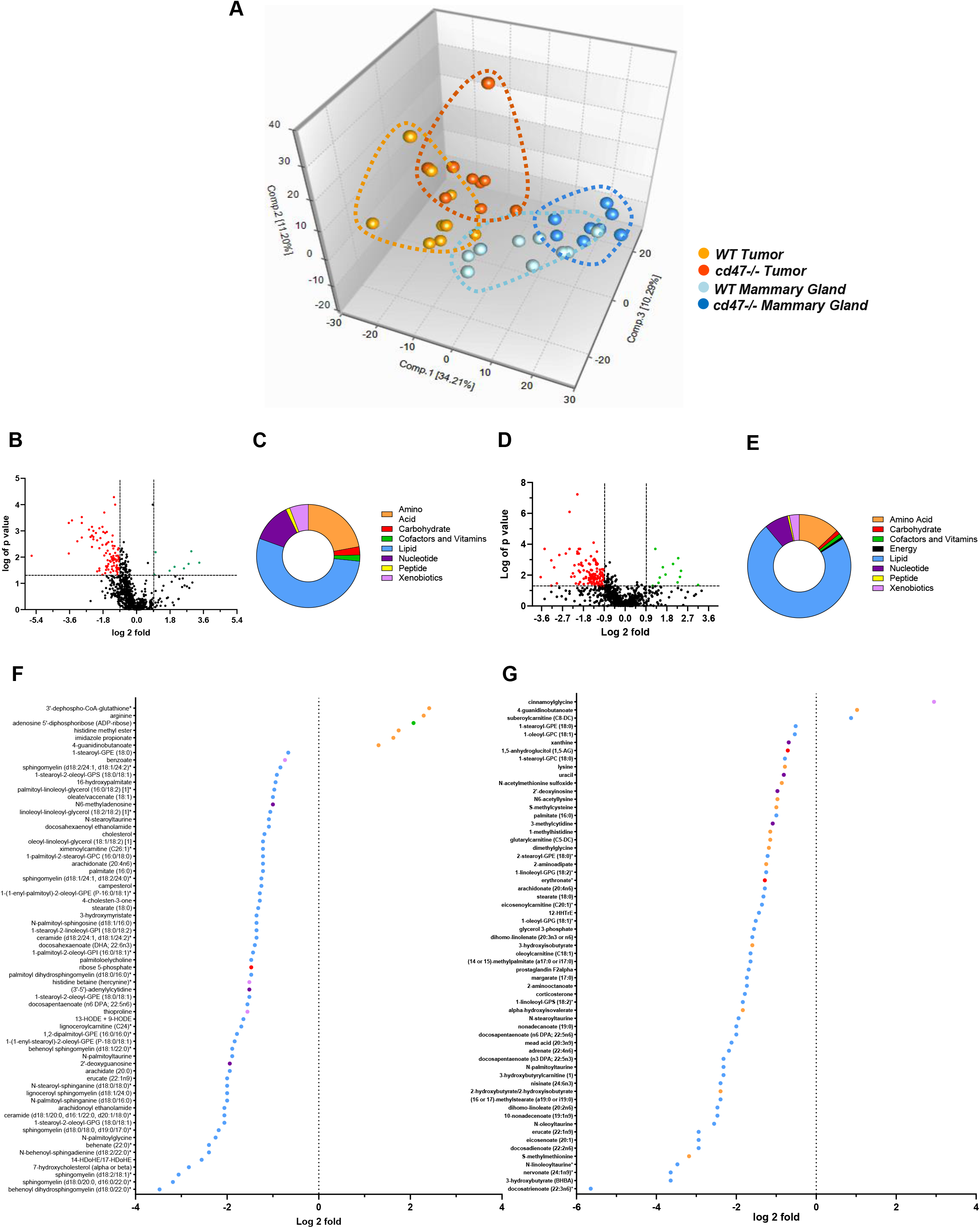
CD47 differentially regulates metabolites in breast tumors and mammary glands. Different biochemical signatures were observed in tumor and mammary gland tissue of WT and *cd47*-/-mice. **(A)** Based on its global metabolomic profiles, a principal component analysis clustered WT and *cd47*-/- breast tumor and mammary gland tissue. It was determined that **(B)** 162 breast tumors and **(D)** 154 mammary gland metabolites were significantly different between WT and *cd47*-/- genotypes (p<0.05). **(C, E)** A pie graph displays the percentage of these statistically significant metabolites based on their respective metabolic pathway. Further analysis displays the **(F)** breast tumor and **(G)** mammary gland metabolites achieving statistical significance of p≤0.01. (n=9/group)

### CD47 expression regulates metabolite markers of proliferation

The proliferation of cancer cells within tumors occurs at a higher rate compared to normal tissue. Metabolites associated with polyamine metabolism, phospholipid synthesis, and membrane phospholipids can indicate how cells proliferate. Polyamines are involved in cell proliferation and division, specifically in cancer cells, as negatively charged DNA can bind to their polycationic surfaces ^22^. A significant increase in polyamines was present in breast tumors compared to mammary glands in both genotypes **(Table I).** Specifically, polyamines like putrescine and spermidine were increased in breast tumors compared to mammary glands in both genotypes **(Figure 3A-B)**.

**Figure 3:**
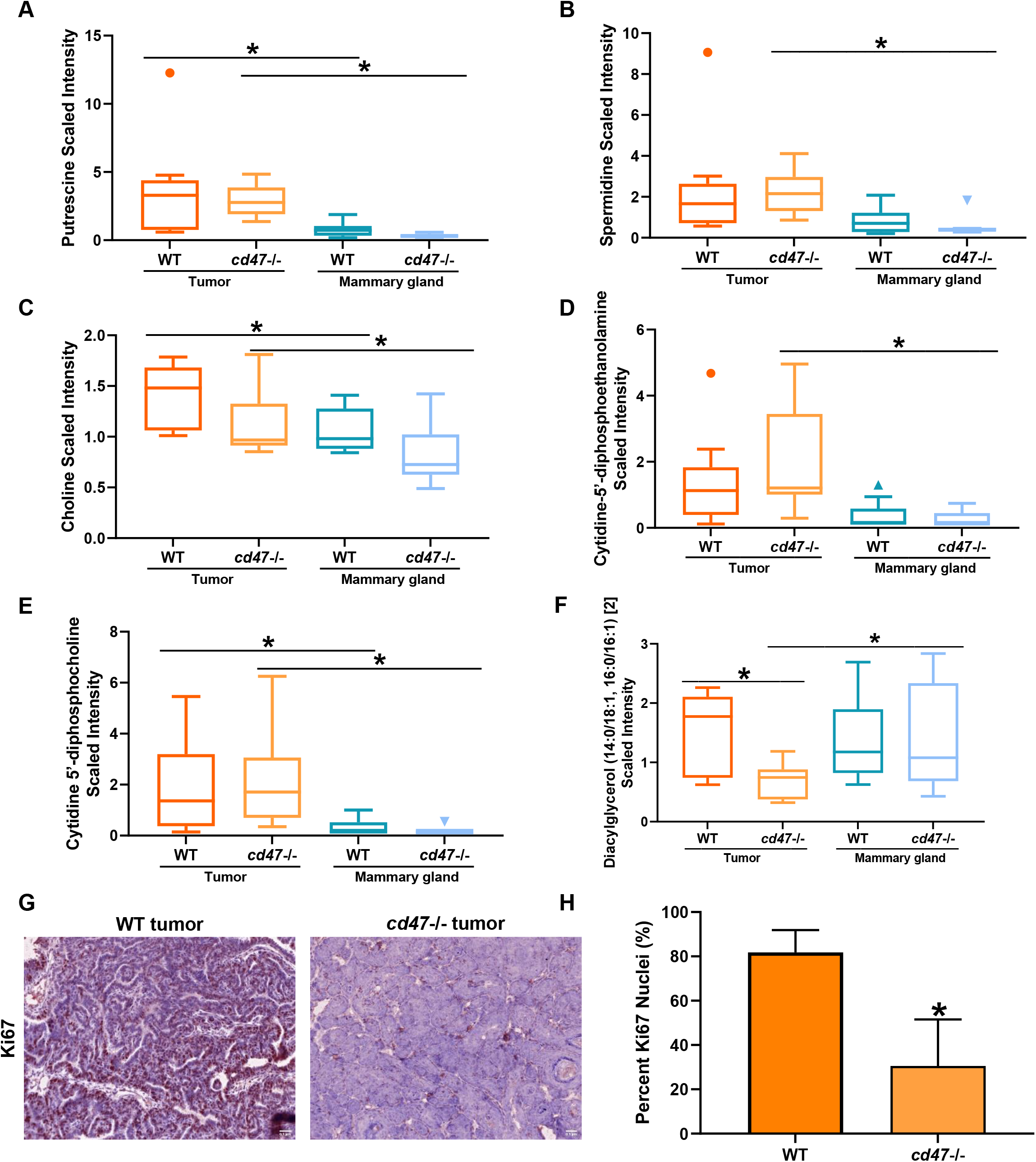
CD47 expression differentially regulates metabolites associated with cell proliferation. Increased metabolites associated with polyamine metabolism, including **(A)** putrescine and **(B)** spermidine in breast tumors compared to mammary glands. Breast tumors have increased metabolites associated with phospholipid metabolism, including **(C)** choline, **(D)** cytidine-5’-diphosphoethanolamine, and **(E)** cytidine 5’-diphosphocholine compared to mammary glands. *cd47*-/- breast tumors have decreased **(F)** diacylglycerol compared to WT breast tumors. (*p<0.05, n=9/group) **(G-H)** Ki67 positive nuclei was decreased in *cd47*-/- breast tumors compared to WT breast tumors (*p<0.05, n=4/group).

**Table I:** Fold changes of metabolites associated with proliferation.

|  |  | Two- way ANOVA Contrasts |  |  |
| --- | --- | --- | --- | --- |
|  |  | Fold of Change |  |  |
|  |  | Tumor Mammary Gland |  | *cd47*<sup>-/-</sup> WT |
|  |  | WT | *cd47*<sup>-/-</sup> | Tumor |
| Polyamine Metabolism | Putrescine | 4.47 | 8.75 |  |
|  | N-acetyl-isoputrescine* | 1.19 | 1.77 |  |
|  | Spermidine | 2.91 | 4.02 |  |
|  | N('1)-acetylspermidine | 5.48 | 21.14 |  |
|  | N1,N12-diacetylspermine | 20.38 | 21.19 |  |
|  | 5-methylthioadenosine (MTA) | 6.85 | 7.47 |  |
|  | N-acetylputrescine | 12.23 | 11.19 |  |
| Phospholipid synthesis | Choline | 1.34 | 1.35 |  |
|  | Choline phosphate | 2.07 | 3.27 |  |
|  | Cytidine 5'-diphosphocholine | 5.64 | 10.71 |  |
|  | Glycerophosphorylcholine (GPC) | 1.66 | 2.04 |  |
|  | Phosphoethanolamine | 1.16 | 1.98 |  |
|  | Cytidine-5'-diphosphoethanolamine | 3.77 | 7.17 |  |
|  | Glycerophosphoethanolamine | 1.86 | 2.45 |  |
|  | Glycerophosphoserine* | 1.01 | 1.39 |  |
|  | Glycerophosphoinositol* | 1.97 | 2.08 |  |
|  | Trimethylamine N-oxide | 1.02 | 1.68 |  |
| Phosphatidylcholine (PC) | 1-myristoyl-2-palmitoyl-GPC (14:0/16:0) | 2.74 | 2.17 |  |
|  | 1-myristoyl-2-arachidonoyl-GPC (14:0/20:4) | 3.96 | 3.65 |  |
|  | 1,2-dipalmitoyl-GPC (16:0/16:0) | 1.97 | 1.59 |  |
|  | 1-palmitoyl-2-stearoyl-GPC (16:0/18:0) | 4.27 | 1.85 |  |
| Phosphatidylethanolamine (PE) | 1,2-dipalmitoyl-GPE (16:0/16:0)* | 10.26 | 2.14 |  |
|  | 1-palmitoyl-2-oleoyl-GPE (16:0/18:1) | 2.45 | 1.37 |  |
| Phosphatidylglycerol (PG) | 1-palmitoyl-2-oleoyl-GPG (16:0/18:1) | 3.50 | 1.92 |  |
|  | 1-palmitoyl-2-linoleoyl-GPG (16:0/18:2) | 1.87 | 1.25 |  |
|  | 1-stearoyl-2-oleoyl-GPG (18:0/18:1) | 6.02 | 1.56 |  |
| Phosphatidylinositol (PI) | 1-palmitoyl-2-oleoyl-GPI (16:0/18:1)* | 8.60 | 2.21 |  |
|  | 1-stearoyl-2-linoleoyl-GPI (18:0/18:2) | 2.39 | 1.02 |  |
|  | 1,2-dioleoyl-GPI (18:1/18:1) | 1.61 | 0.65 |  |
|  | 1-stearoyl-2-arachidonoyl-GPI (18:0/20:4) | 1.48 | 0.99 |  |
| Diacylglycerols | Diacylglycerol (14:0/18:1, 16:0/16:1) [2] |  |  | 0.62 |
|  | Palmitoyl-oleoyl-glycerol (16:0/18:1) [1] |  |  | 0.41 |
|  | Palmitoyl-oleoyl-glycerol (16:0/18:1) [2] |  |  | 0.44 |
|  | Palmitoyl-linoleoyl-glycerol (16:0/18:2) [1] |  |  | 0.51 |
|  | Palmitoyl-linoleoyl-glycerol (16:0/18:2) [2] |  |  | 0.49 |
|  | Palmitoleoyl-linoleoyl-glycerol (16:1/18:2) [1] |  |  | 0.54 |
|  | Palmitoyl-dihomo-linolenoyl-glycerol (16:0/20:3n3 or 6) [2] |  |  | 0.65 |
|  | Palmitoyl-arachidonoyl-glycerol (16:0/20:4) [1] |  |  | 0.69 |
|  | Palmitoyl-arachidonoyl-glycerol (16:0/20:4) [2] |  |  | 0.57 |
|  | Palmitoyl-docosaheptaenoyl-glycerol (16:0/22:6) [1] |  |  | 0.87 |
|  | Stearoyl-linoleoyl-glycerol (18:0/18:2) [2] |  |  | 0.47 |
|  | Oleoyl-oleoyl-glycerol (18:1/18:1) [1] |  |  | 0.45 |
|  | Oleoyl-oleoyl-glycerol (18:1/18:1) [2] |  |  | 0.39 |
|  | Oleoyl-linoleoyl-glycerol (18:1/18:2) [1] |  |  | 0.43 |
|  | Oleoyl-linoleoyl-glycerol (18:1/18:2) [2] |  |  | 0.41 |
|  | Linoleoyl-linoleoyl-glycerol (18:2/18:2) [1] |  |  | 0.48 |
|  | Linoleoyl-linoleoyl-glycerol (18:2/18:2) [2] |  |  | 0.41 |
|  | Linoleoyl-linolenoyl-glycerol (18:2/18:3) [1] |  |  | 0.46 |
|  | Oleoyl-arachidonoyl-glycerol (18:1/20:4) [2] |  |  | 0.58 |
|  | Linoleoyl-arachidonoyl-glycerol (18:2/20:4) [2] |  |  | 0.59 |
|  | Stearoyl-docosaheptaenoyl-glycerol (18:0/22:6) [1] |  |  | 0.66 |
|  | Stearoyl-docosaheptaenoyl-glycerol (18:0/22:6) [2] |  |  | 0.57 |

On the other hand, the largest cell membrane component is phospholipids to support cell growth and survival ^23^. Therefore, we examined metabolites associated with phospholipid synthesis and membrane phospholipids as an increase in these respective metabolites indicates increased cell proliferation. A significant increase in metabolites related to phospholipid synthesis occurred in both genotypes of breast tumors compared to mammary glands **(Table I)**. Specifically, choline, cytidine-5’-diphosphoethanolamine, and cytidine 5’-diphosphocholine were increased in breast tumor tissue compared to mammary gland tissue **(Figure 3C-E).** In breast tumors, a significant increase in metabolites associated with membrane phospholipids like phosphatidylcholines, phosphatidylethanolamine, phosphatidylglycerol, and phosphatidylinositol was also increased over mammary glands in both WT and *cd47*-/- mice **(Table I)**.

Although a significant increase in phospholipid synthesis was observed in both WT and *cd47*-/- breast tumors, changes in diacylglycerols were observed between genotypes of breast tumors. A significant decrease in diacylglycerols was observed between *cd47*-/- breast tumors compared to WT breast tumors **(Figure 3F**, **Table I)**. This may indicate reduced proliferation in tumors due to the lack of CD47 receptor expression. Therefore, to determine how CD47 expression impacts proliferation, we stained breast tumor sections for Ki67, a proliferation marker. We observed a significant decrease in Ki67 positive cells within *cd47*-/- breast tumors compared to WT breast tumors **(Figure 3G-H)**. This was further validated through a spatial proteomic analysis where *cd47*-/- breast tumors significantly decreased Ki67 protein expression compared to WT breast tumors **(Supplementary Figure 1E)**.

### CD47 preferentially downregulates fatty acid metabolism in breast tumors

The tumor microenvironment can play a role in the metabolic processes occurring within cancer cells and normal surrounding tissue. Breast tissue is rich in adipocytes which are responsible for storing fat. Therefore, tumors developing within breast tissue have increased lipid precursors readily available to undergo fatty acid oxidation and progress disease ^24, 25^. We examined metabolites associated with fatty acid oxidation and how CD47 expression impacts this metabolic process. Several long-chained fatty acids and metabolites associated with polyunsaturated fatty acid metabolism were reduced in breast tumors compared to mammary glands **(Table II)**. Of note, myristate and palmitoleate, which are long-chain fatty acids, and hexadecadienoate and linoleate, metabolites associated with polyunsaturated fatty acid metabolism, were decreased in both breast tumors compared to mammary glands of both genotypes **(Figure 4A-D)**. However, an increase in several carnitine-conjugated fatty acids, like arachidoylcarnitine and myristoylcarnitine, was observed in breast tumors compared to mammary glands **(Figure 4E-F**, **Table II)**. These fatty acids are synthesized from excess acetyl-CoA produced by mitochondrial β-oxidation ^26^. While breast tumors exhibited elevated metabolites associated with the citric acid cycle compared to mammary glands **(Supplementary Figure 3A-E),** minimal differences were observed between metabolites of the citric acid cycle between WT and *cd47*-/- tissues, suggesting CD47 expression has little effect on the citric acid cycle **(Supplementary Figure 3 A-E, Supplementary Table I)**. Thus, suggesting that CD47 signaling favors energy production from fatty acid metabolism in the breast tumor microenvironment.

**Figure 4:**
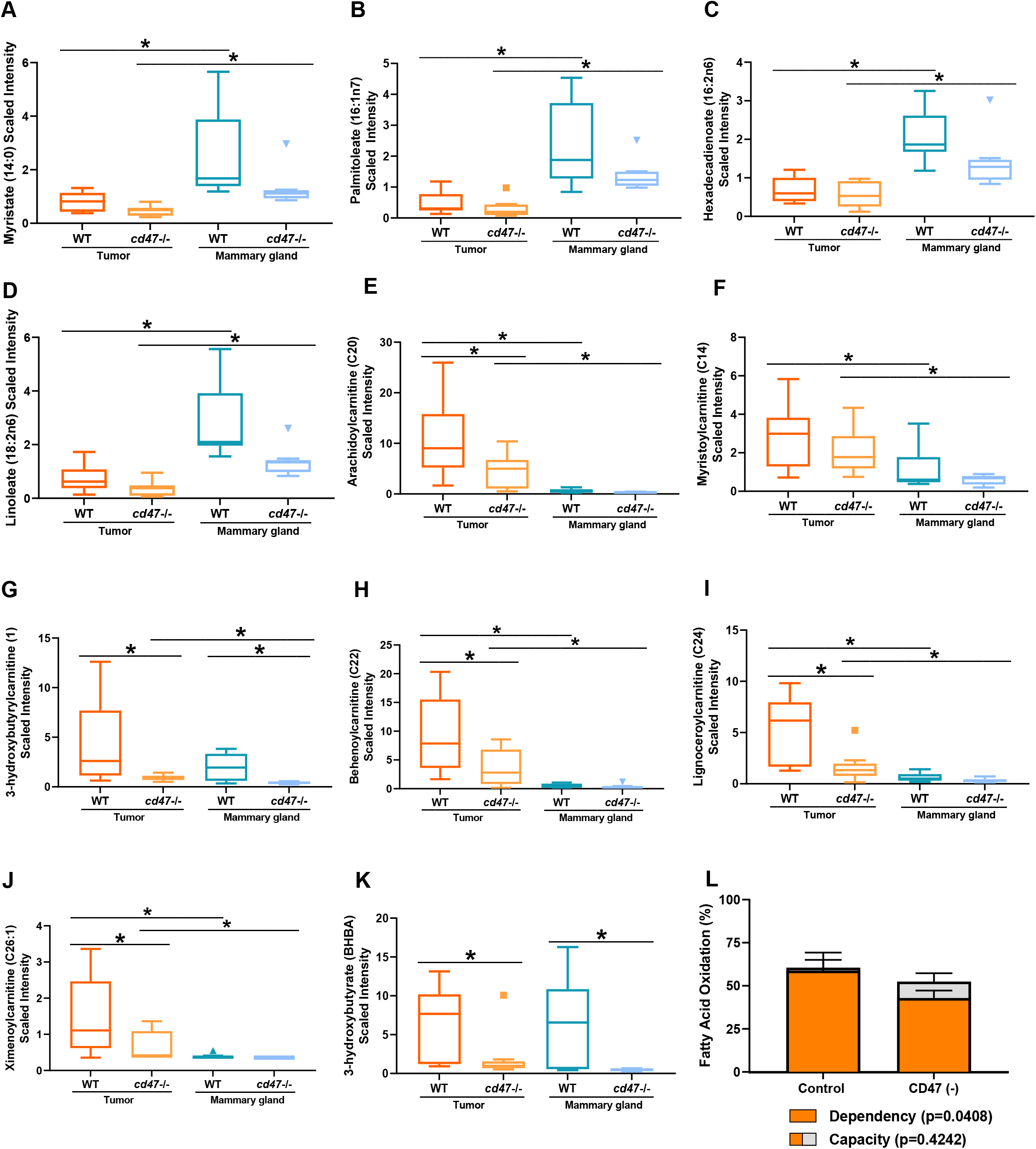
Fatty acid metabolism is mediated by CD47 expression. There is a significant decrease in long-chained fatty acids, specifically **(A)** myristate and **(B)** palmitoleate, in breast tumors compared to mammary glands. Metabolites associated with polyunsaturated fatty acid metabolism like **(C)** hexadecadienoate and **(D)** linoleate were significantly decreased in breast tumor tissue compared to mammary glands. An increase in carnitine-conjugated fatty acids like **(E)** arachidoylcarnitine and **(F)** myristoylcarnitine was observed in breast tumors compared to mammary glands. Both *cd47*-/- breast tumors and mammary glands had a significant decrease in metabolites associated with fatty acid metabolism like **(G)** 3-hydroxy butyryl carnitine **(H)** behenoylcarnitine, **(I)** lignoceroylcarnitine and **(J)** ximenoylcarnitine, and ketone body **(K)** 3-hydroxybutyrate (BHBA). (*p<0.05, n=9/group) **(L)** CD47 targeted EMT-6 cells have a decrease in fatty acid dependency compared to WT EMT-6 cells (*p<0.05, n=3).

**Table II:** Fold change between metabolites associated with polyunsaturated fatty acid metabolism, long-chain fatty acids, and fatty acid metabolism.

|  |  | Two-way ANOVA Contrasts |  |  |  |
| --- | --- | --- | --- | --- | --- |
|  |  | Tumor |  | cd47<sup>-/-</sup> |  |
|  |  | Mammary Gland |  | WT |  |
|  |  | WT | cd47 null | Tumor | Mammary Gland |
| Polyunsaturated Fatty Acid Metabolism | Docosapentaenoate (n3 DPA; 22:5n3) | 0.56 | 1.14 | 0.41 | 0.20 |
|  | Docosahexaenoate (DHA; 22:6n3) | 1.20 | 1.13 | 0.38 | 0.40 |
|  | Nisinate (24:6n3) | 1.11 | 2.04 | 0.36 | 0.19 |
|  | Linoleate (18:2n6) | 0.26 | 0.26 | 0.49 | 0.48 |
|  | Linolenate α or γ (18:3n3 or 6) | 0.14 | 0.16 | 0.75 | 0.66 |
|  | Dihomo-linolenate (20:3n3 or n6) | 0.93 | 1.15 | 0.40 | 0.33 |
|  | Arachidonate (20:4n6) | 1.56 | 1.63 | 0.43 | 0.41 |
| Long Chain Fatty Acid | Myristate (14:0) | 0.31 | 0.36 | 0.59 | 0.51 |
|  | Palmitate (16:0) | 0.72 | 0.61 | 0.43 | 0.50 |
|  | Palmitoleate (16:1n7) | 0.21 | 0.24 | 0.67 | 0.58 |
|  | Stearate (18:0) | 1.21 | 1.23 | 0.40 | 0.40 |
|  | Oleate/vaccenate (18:1) | 0.34 | 0.39 | 0.50 | 0.43 |
|  | Nonadecanoate (19:0) | 0.60 | 1.13 | 0.48 | 0.25 |
|  | 10-nonadecenoate (19:1n9) | 0.17 | 0.55 | 0.59 | 0.18 |
| Fatty Acid Metabolism (Acyl Carnitine) | Acetylcarnitine (C2) | 1.26 | 1.33 | 0.79 | 0.75 |
|  | 3-hydroxybutyrylcarnitine (1) | 2.10 | 3.15 | 0.31 | 0.20 |
|  | Hexanoylcarnitine (C6) | 1.61 | 1.81 | 0.92 | 0.82 |
|  | 3-hydroxyhexanoylcarnitine (2) | 1.49 | 2.11 | 0.75 | 0.53 |
|  | 3-hydroxyhexanoylcarnitine (1) | 1.80 | 2.01 | 0.68 | 0.61 |
|  | Octanoylcarnitine (C8) | 3.92 | 4.02 | 0.87 | 0.84 |
|  | Decanoylcarnitine (C10) | 6.88 | 4.81 | 0.71 | 1.01 |
|  | 5-dodecenoylcarnitine (C12:1) | 3.73 | 2.66 | 0.64 | 0.89 |
|  | Laurylcarnitine (C12) | 1.71 | 2.59 | 0.75 | 0.49 |
|  | Myristoylcarnitine (C14) | 2.37 | 3.64 | 0.76 | 0.50 |
|  | Palmitoylcarnitine (C16) | 3.28 | 4.27 | 0.68 | 0.52 |
|  | Palmitoleoylcarnitine (C16:1)* | 1.60 | 2.59 | 0.79 | 0.48 |
|  | Stearoylcarnitine (C18) | 5.87 | 8.49 | 0.58 | 0.40 |
|  | Linoleoylcarnitine (C18:2)* | 1.53 | 2.91 | 0.76 | 0.40 |
|  | 3-hydroxyoleoylcarnitine | 1.56 | 2.08 | 0.63 | 0.47 |
|  | Oleoylcarnitine (C18:1) | 1.30 | 2.95 | 0.73 | 0.32 |
|  | Suberoylcarnitine (C8-DC) | 0.53 | 0.29 | 1.00 | 1.83 |
|  | Octadecenedioylcarnitine (C18:1-DC)* | 1.62 | 1.52 | 1.18 | 1.25 |
|  | Arachidoylcarnitine (C20)* | 18.35 | 16.61 | 0.43 | 0.47 |
|  | Adrenoylcarnitine (C22:4)* | 13.40 | 16.81 | 0.74 | 0.59 |
|  | Behenoylcarnitine (C22)* | 17.86 | 10.27 | 0.38 | 0.66 |
|  | Dihomo-linolenoylcarnitine (C20:3n3 or 6)* | 6.53 | 10.40 | 0.81 | 0.51 |
|  | Eicosenoylcarnitine (C20:1)* | 3.13 | 5.26 | 0.65 | 0.39 |
|  | Erucoylcarnitine (C22:1)* | 7.85 | 14.55 | 0.70 | 0.38 |
|  | Docosatrienoylcarnitine (C22:3)* | 9.43 | 10.09 | 0.73 | 0.68 |
|  | Docosapentaenoylcarnitine (C22:5n3)* | 26.56 | 11.34 | 0.54 | 1.26 |
|  | Docosahexaenoylcarnitine (C22:6)* | 10.45 | 3.82 | 0.43 | 1.19 |
|  | Lignoceroylcarnitine (C24)* | 8.50 | 4.72 | 0.31 | 0.56 |
|  | Margaroylcarnitine (C17)* | 2.49 | 5.41 | 0.93 | 0.43 |
|  | Nervonoylcarnitine (C24:1)* | 8.33 | 10.28 | 0.57 | 0.46 |
|  | Ximenoylcarnitine (C26:1)* | 3.96 | 1.83 | 0.43 | 0.93 |
|  | 3-hydroxypalmitoylcarnitine | 2.75 | 2.95 | 0.63 | 0.59 |

Furthermore, our hierarchical clustering analysis shows a global decrease in fatty acid metabolites in *cd47*-/- breast tumors compared to WT breast tumors **(Supplementary Figure 2)**. Additionally, *cd47*-/- breast tumors were significantly decreased in carnitine conjugated fatty acids like arachidoylcarnitine, 3-hydroxy butyryl carnitine, behenoylcarnitine, lignoceroylcarnitine, and ximenoylcarnitine compared to WT breast tumors **(Figure 4E, 4G-J).** Ketone body 3- hydroxybutyrate (BHBA) was also observed to be decreased in *cd47*-/- mice compared to WT mice **(Figure 4K)**. A decrease in these metabolites suggests a reduction in fatty acid availability and metabolism due to the lack of CD47 expression. Cancer cells undergo aerobic glycolysis, more commonly known as the Warburg effect, to produce energy, resulting in increased glucose consumption and end-products like pyruvate and lactate ^27^. Glycolysis was activated in breast tumors compared to mammary glands in both genotypes since a decrease in glucose and an increase in pyruvate and lactate were observed **(Supplementary Figures 3G-H, Supplementary Table II).** Therefore, even though breast tumor growth may follow Warburg metabolism, our data shows that absence of CD47 may result in the inhibition of pathways that result in the downregulation of fatty acid metabolism. To determine whether this may be due to the circumstances of the breast tumor microenvironment or due to an autonomous regulation of fatty acid metabolism by CD47, we performed a Seahorse Mito Fuel Flex Test assay to examine metabolic pathway dependency for mitochondrial oxidation of EMT-6 cells based on CD47 expression. The dependency on fatty acid metabolism is calculated by considering the mitochondrial function measured by OCR in the presence or absence of etomoxir (Fatty acid oxidation inhibitor). Targeting CD47 on EMT-6 cells decreased the dependency of fatty acid oxidation compared to control EMT-6 cells, validating the metabolomic analysis **(Figure 4L)**. Also consistent with our metabolomics analysis, the absence of CD47 did not regulate a glycolytic pathway in tumors (**Supplementary Figures 3I-3L**). Thus, the lack of CD47 may directly control fatty acid oxidation to reduce tumor burden.

### CD47 modulates inflammation within breast tumors

Eicosanoids are lipid molecules with signaling and pro-inflammatory functions as they are bioactive derivatives of omega-6 PUFA arachidonate ^28^. When dysregulation of eicosanoids occurs, as in cancer, they can induce a state of inflammation ^29^. Our metabolomics data shows a significant decrease in eicosanoids, including prostaglandin F2α and 12- Hydroxyheptadecatrienoic acid (12-HHTre) in *cd47*-/- tissue compared to WT tissue **(Figure 5A-B**, **Table III)**. This suggests that decreased eicosanoids may decrease the inflammatory state within *cd47*-/- tissue compared to WT tissue.

**Figure 5:**
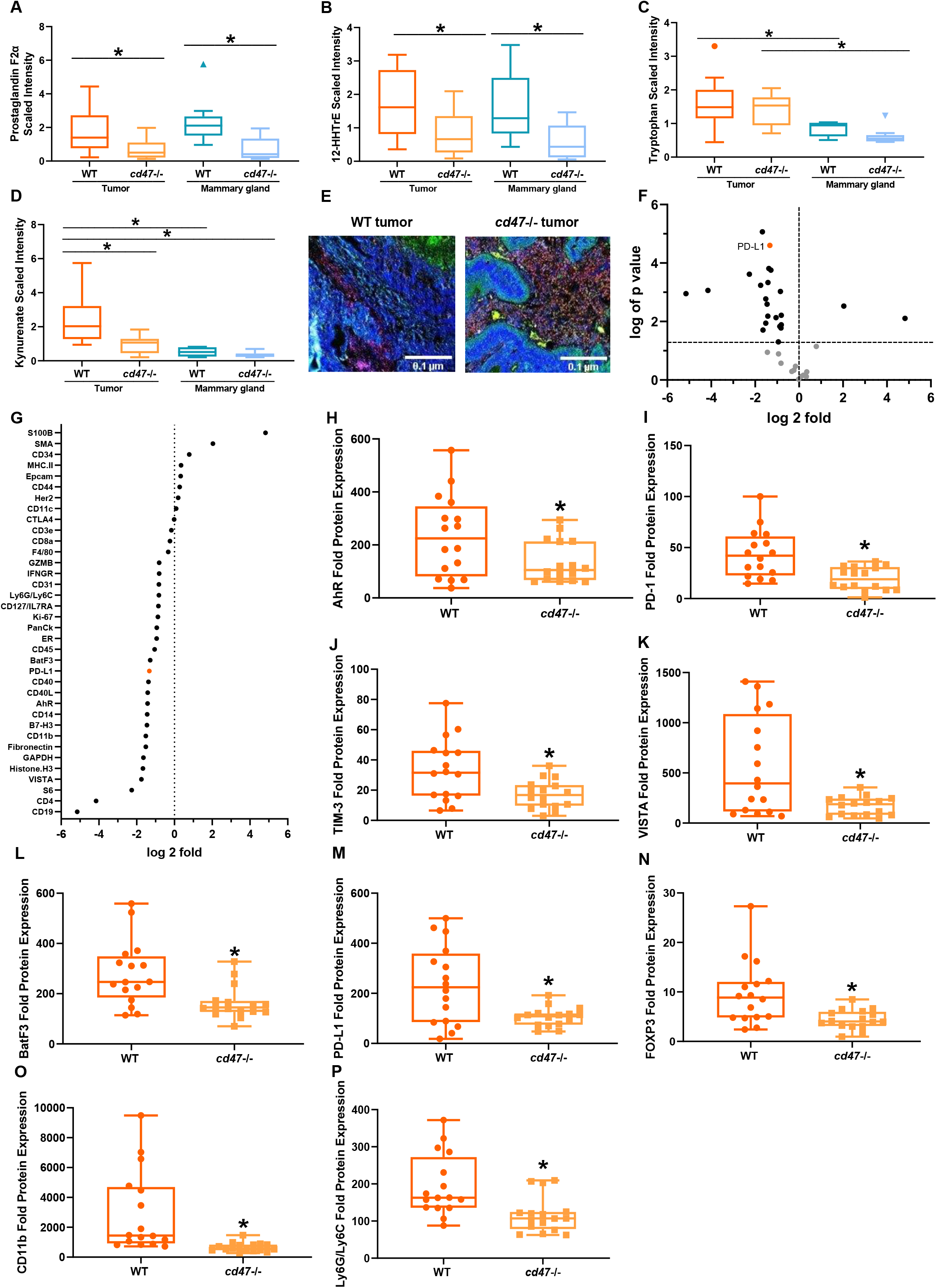
CD47 mediates intratumoral inflammation. Eicosanoids are decreased in *cd47*-/- tissue compared to WT tissue, specifically **(A)** prostaglandin F2α and **(B)** 12- HHTre. **(C)** Tryptophan metabolites are increased in tumor tissue compared to the mammary gland of both genotypes. **(D)** Kynurenate is increased in WT breast tumors compared to the mammary gland and *cd47*-/- breast tumors. (*p<0.05, n=9/group) **(E)** Fluorescent antibodies for lymphocytes (CD45+, red), T cells (CD3+, yellow), cancer cells (PanCK, green), and nuclei (DAPI, blue) stained the tumor sections to determine regions of interest to undergo spatial proteomic analysis. 36 proteins were analyzed through DSP where **(F-G)** 2 proteins were enriched in the *cd47*-/- breast tumors while 21 were enriched in the WT breast tumors. **(H)** AhR receptor expression is decreased in *cd47*-/- breast tumors compared to WT breast tumors. WT breast tumors have an increase in immunosuppressive receptors like **(I)** PD-1, **(J)** TIM-3, **(K)** VISTA, **(L)** BatF3, and **(M)** PD- L1 as well as an increase in **(N)** FOXP3 **(O)** CD11b+ and **(P)** Ly6G/Ly6C+ immunosuppressive immune cells compared to *cd47*-/- breast tumors. (*p<0.05, n=4/group)

**Table III:**
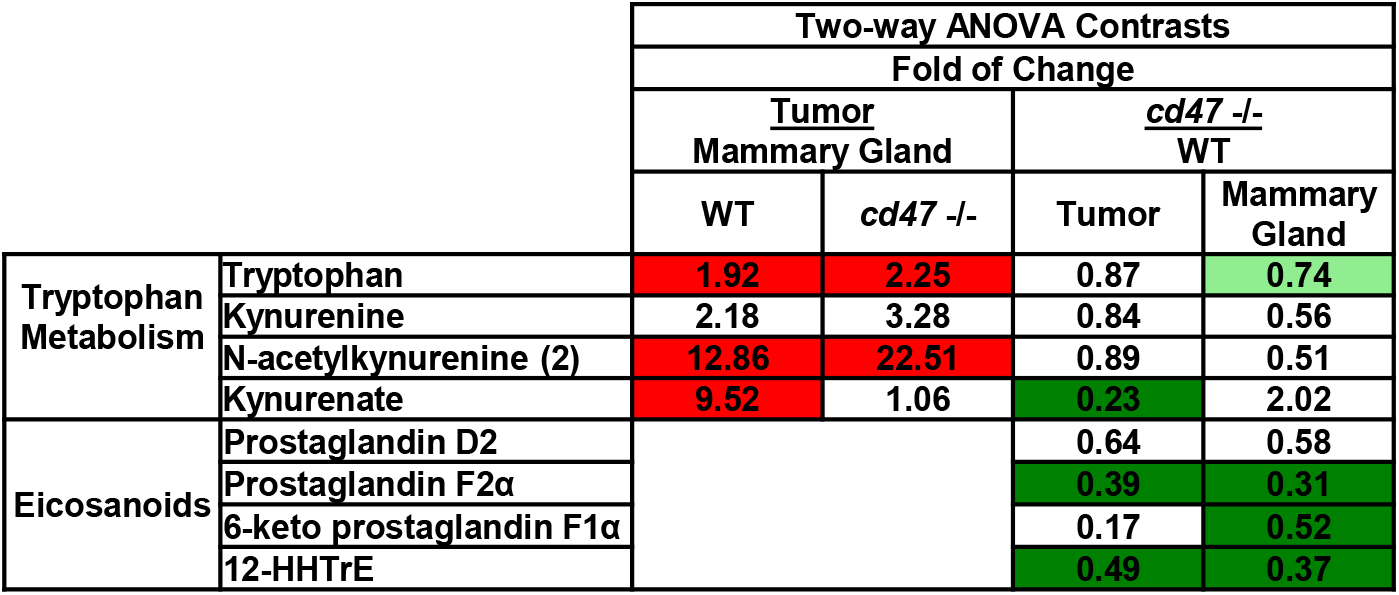
Fold change of metabolites associated with inflammation.

The inflammatory state within malignant tissue can also create an environment to support tumorigenesis compared to normal tissue ^30^. Tryptophan metabolites can be a reflection of an altered inflammatory state. IDO is activated by pro-inflammatory cytokines like interferon-gamma (IFNγ) and tumor necrosis factor-alpha (TNFα) to catalyze the conversion of tryptophan to kynurenine ^31^. A significant increase in tryptophan was observed in breast tumors compared to mammary glands in both genotypes **(Figure 5C**, **Table III).** Additionally, a significant increase in kynurenate, a product of tryptophan metabolism, was observed in WT breast tumors compared to mammary glands; however, this metabolite was significantly decreased in *cd47*-/- breast tumors compared to WT breast tumors **(Figure 5D**, **Table III)**. Reactive oxygen species can promote kynurenate production ^32^. Previous data has shown that *cd47*-/- lung tissue and T cells have a greater capacity to handle radiation-induced oxidative stress ^15, 16^. Our data support these findings as *cd47*-/- breast tumors have increased oxidized and reduced glutathione (GSSG and GSH) compared to WT breast tumors **(Supplementary Table III)**, thus suggesting a possible mechanism for stimulation of inflammation in WT tumor tissue.

Kynurenate acts as a ligand to the aryl hydrocarbon receptor (AhR), mediating immunosuppression ^33^. When kynurenate binds to AhR, it dimerizes with AhR nuclear translocator to function as a transcription factor and induce an immunosuppressive environment ^33^. We examined the overall protein expression of several proteins associated with immunosuppressive signaling within breast tumor tissue through spatial proteomic analysis **(Figure 5G)**. Of the 36 proteins analyzed, 2 were enriched in the *cd47*-/- breast tumors, while 21 were enriched in the WT breast tumors **(Figure 5F).** We observed a significant decrease in AhR receptor expression within *cd47*-/- breast tumors compared to WT breast tumors, suggesting a decreased inflammatory state mediated by CD47 expression **(Figure 5H)**. Other metabolites like itaconate and BHBA, which support inflammation, were also reduced in *cd47*-/- tissue compared to WT tissue **(Supplementary Table I and Figure 4K).** Therefore, it is evident that CD47 regulates inflammation, supporting previous studies where *cd47*-/- adipose tissue had a reduction in inflammation, resulting in a decreased accumulation of pro-inflammatory M2-like macrophages **(Supplementary Figure 4A-C)** ^34, 35^.

### CD47 expression mediates immunosuppression signaling within the breast tumor microenvironment

Since inflammation can impair antitumor immune response, we further examined immunosuppressive receptors and cell types within WT and *cd47*-/- breast tumors through spatial proteomic analysis. An increase in T cell expressing immune checkpoint proteins like programmed death protein 1 (PD-1), T cell immunoglobulin and mucin-domain containing-3 (TIM- 3), V-domain Ig suppressor of T cell activation (VISTA), basic leucine zipper transcription factor ATF-like 3 (BatF3) was observed in WT breast tumors compared *cd47*-/- breast tumors **(Figure 5I-L)**. Additionally, PD-L1, an immune checkpoint protein located on cancer cells, was increased within WT breast tumors compared to *cd47*-/- breast tumors **(Figure 5M)**. Forkhead box protein P3 (FOXP3), a marker for T regulatory cells, and CD11b+ and Ly6c+, markers of myeloid-derived suppressing cells, were also increased in WT breast tumors compared to *cd47*-/- breast tumors **(Figure 5N-P)**.

### Targeting CD47 sensitizes TNBC tumors to immune checkpoint blockade therapy

Immune checkpoint blockade therapies targeting PD-L1 are FDA approved to treat metastatic TNBC; however, therapeutic response is limited ^3, 36^. Additionally, as discussed earlier, WT breast tumors have elevated expression of immunosuppressive proteins, like PD-L1, compared to *cd47*-/- breast tumors. Therefore, we examined how targeting CD47 would impact the tumor burden of mice receiving anti-PD-L1 treatment in an EMT-6 mouse model **(Figure 6A)**. Results were compared to control as no significant difference occurred in tumor volume of IgG and CTRLM treated mice **(Supplementary Figure 4E)**. A decrease in tumor volume was observed by targeting CD47 and PD-L1 as monotherapies **(Figure 6B)**. When these treatments were received in combination, a further reduction in tumor volume was observed compared to anti-PD-L1 treatment and control **(Figure 6B)**. This decrease in tumor burden may be mediated by infiltrating CD8+ T cells with enhanced antitumor function as using these therapies in combination increased granzyme B secreting intratumoral CD8+ T cells compared to monotherapies and control **(Figure 6C-E)**.

**Figure 6:**
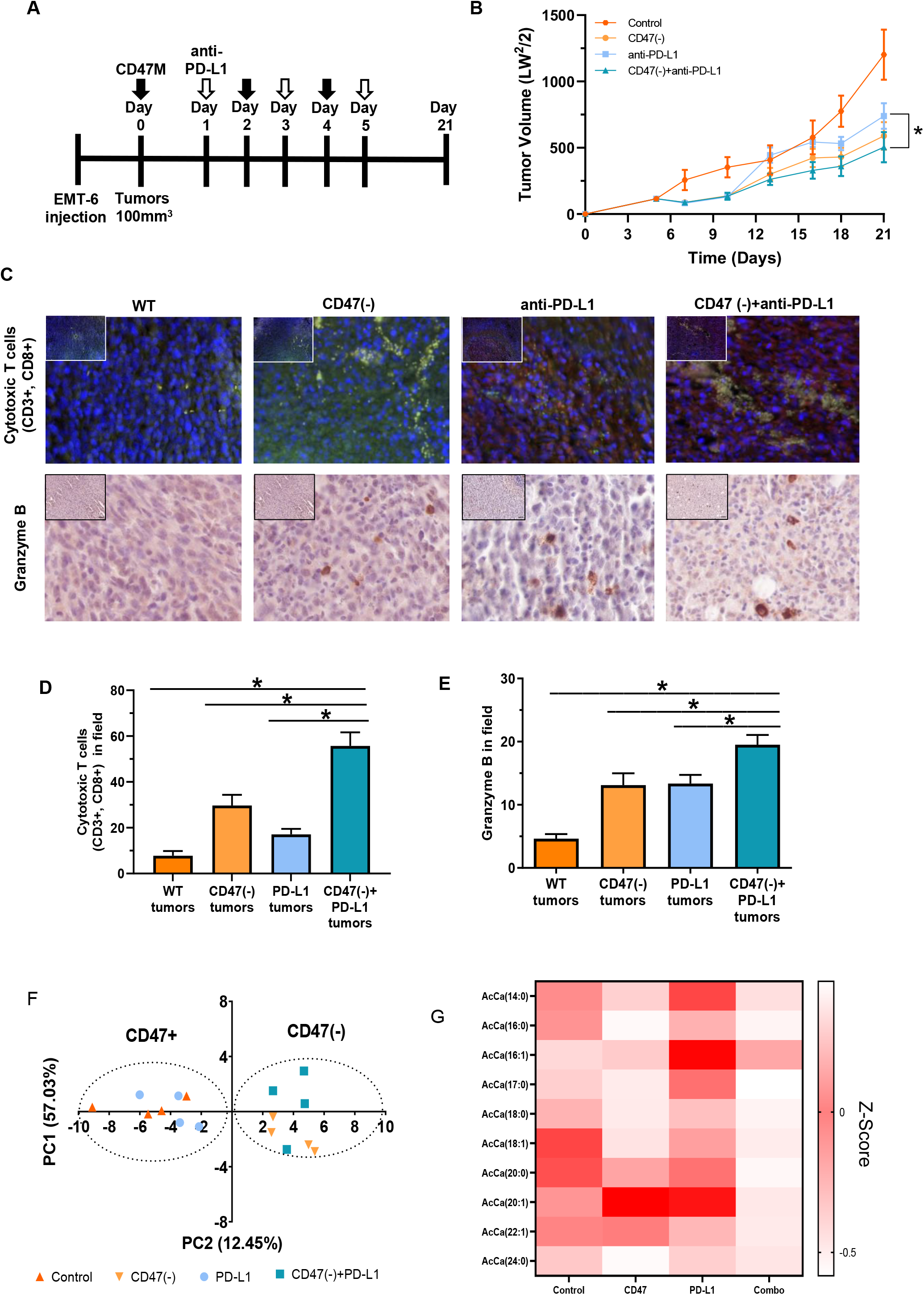
CD47 blockade sensitizes tumors to immune checkpoint blockade therapy. **(A)** Schematic of orthotopic TNBC mouse model treatment regimen. At week 6- 8 of age, female Balb/c mice were injected orthotopically into the mammary fat pad with EMT-6 TNBC cells. Mice received alternating day treatments of CD47M and anti-PD-L1 over 6 days. Mice were euthanized on day 21 or when tumors reached 1500mm^3^. **(B)** Targeting CD47 in combination with anti-PD-L1 decreased tumor volume compared to monotherapy treatments and controls. **(C-D)** An increase in intratumoral CD8+ T cells (CD3+ (red), CD8+ (green), DAPI (blue)) and **(C, E)** increased granzyme B release was observed in combination-treated breast tumors compared to monotherapy and control. (*p<0.05, n=4-8). (F) Principal component of EMT-6 tumor model lipidomic analysis and (G) Heat map of Z-scores of acyl-carnitines detected (n=4/group).

Since we observed a depletion of lipid metabolites in our DMBA model, we subjected tumors from the EMT-6 mouse model to lipidomic analysis to determine whether targeting CD47 in the presence or absences of PD-L1 would impact lipid metabolism. Our principal component analysis shows separation of samples based on presence or depletion of CD47 using antisense morpholino (CD47+/CD47(-) respectively, **Figure 6F).** We noted that several acyl carnitines (8 out of 10 detected were down regulated in the absence of CD47 or combination with anti-PD-L1 (**Figure 6G)**. This suggests that targeting CD47 also reduces lipid metabolism to potentially enhance response to immune checkpoint blockade.

## Discussion

CD47 is a cell surface receptor that modulates innate and adaptive immune responses in the tumor microenvironment ^4^. Our data showed that CD47 was expressed in primary and matched metastatic tumor tissue, making it a potential therapeutic target for advanced breast cancer. However, aside from immune-regulation, CD47 is implicated in autonomous cell death and survival pathways ^14, 37–39^. Our group and others have shown that blockade of this receptor is protective against stress preserving cell and tissue viability through the regulation of metabolism and metabolism-dependent pathways such as autophagy ^13, 15, 16^. We have previously shown regulation of metabolism due to CD47 receptor expression in different cell and tissue types exposed to ionizing radiation ^15, 16^. These studies have provided insight into radioprotection due to alterations in lung tissue and T cell metabolism mediated by CD47 expression. In a rat model of metabolic syndrome, the absence of CD47 was shown to protect mice from age-related obesity and glucose intolerance ^40^. Bulk RNA sequencing suggested the regulation of several metabolic pathways in adipose tissue, suggesting a prominent role of CD47 in the regulation of bioenergetics ^40^. Therefore, given the interest in incorporating CD47 targeting drugs in clinical oncology, it is critical to understand whether the absence of CD47 in cancer cells impacts tumor metabolism and its implications in the tumor microenvironment.

To determine the effect of CD47 deficiency in tumor metabolism, we subjected tumor tissues harvested from the carcinogen-induced model to global metabolomics analysis. We also subjected mammary glands to metabolic analysis to tease out metabolic changes between normal adjacent tissue to tumor tissue. Robust differences in metabolites were apparent between the breast tumors and mammary glands irrespective of the genotype. These metabolic distinctions were evident in the principal component analysis and consistent with our previous observations that absence of CD47 regulates the metabolism of normal tissue under stress ^16^. Therefore, this is one of the first indications that CD47 regulates tumor metabolism. Thus, we further examined regulated metabolites associated with proliferation, metabolism, inflammation, and immunosuppression modified by each genotype.

Uncontrolled and chronic proliferation is a common characteristic of cancer ^41^. Major differences were observed in metabolites associated with proliferation between tissues and genotypes. Polyamines provide a binding site for DNA to subsequently enhance cell proliferation ^22^. Within cancer, polyamine metabolism is often dysregulated due to the interplay with oncogenes and mutated tumor suppressors ^42–44^. This results in the upregulation of polyamines to enhance cancer cell proliferation to support disease progression. We observed that metabolites associated with the metabolism of polyamines were increased in breast tumors compared to mammary glands in both genotypes. Within polyamine metabolism, putrescine is initially produced by ornithine decarboxylase ^45^. Putrescine is then transformed into other polyamines like spermidine with spermidine synthase ^46^. We observed an increase in both putrescine and spermidine within WT and *cd47*-/- breast tumors compared to mammary glands suggesting a differential regulation between tumor and normal adjacent mammary gland.

Phospholipids also support proliferation as they are the largest component of cell membranes. Elevated phospholipid content has been previously observed in breast tumors compared to adjacent, non-malignant mammary glands ^47^. We observed an increase in metabolites associated with phospholipid synthesis within breast tumors compared to mammary glands. Specifically, metabolites associated with the phospholipid components like phosphatidylcholines, phosphatidylethanolamines, phosphatidylglycerol, and phosphatidylinositol were increased in breast tumors compared to the mammary gland. This data supports previous studies of increased phosphatidylcholines and phosphatidylethanolamines in breast tumors ^48^. It is evident that breast tumors have an increased demand to proliferate and expand compared to mammary glands due to the increase in metabolites associated with polyamine metabolism, phospholipid synthesis, and membrane phospholipids; however, this is not dependent on CD47 receptor expression. Alternatively, a reduction in metabolites related to diacylglycerols was observed in *cd47*-/- breast tumors compared to WT breast tumors. Diacylglycerols act as messengers to induce signaling for cell division by activating protein kinase C ^49^. Therefore, a decrease in metabolites associated with diacylglycerols may indicate a reduction in cell proliferation with tumors lacking CD47 expression. This was validated due to the reduced Ki67, a marker of cell proliferation, expression within *cd47*-/- breast tumors compared to WT breast tumors. Overall, CD47 expression modulates cancer cell proliferation and potentially tumor burden *in vivo* by mediating diacylglycerol metabolites.

When examining the metabolism of breast cancer, it is essential to consider the tumor microenvironment as breast tissue consists of adipocytes and is rich in several precursors for fatty acid metabolism. Elevated rates of fatty acid oxidation occurred in breast tumors compared to mammary glands due to the reduction of long-chained fatty acids and metabolites associated with polyunsaturated fatty acid metabolism. Carnitine-conjugated fatty acids, synthesized from excess acetyl-CoA produced by mitochondrial β-oxidation, were also increased in breast tumors compared to mammary glands, supporting the increased rate of fatty acid metabolism. Still, most long-chain fatty acid metabolites were downregulated in *cd47*-/- breast tumors compared to WT breast tumors. This is consistent with previous reports that reducing fatty acid oxidation by inhibiting the unfolded protein response is associated with reducing CD47 expression and enhanced macrophage infiltration ^50^. Although breast tumors increase fatty acid metabolism compared to mammary glands, there was differential regulation between genotypes. *cd47*-/- breast tumors significantly decreased BHBA and several carnitines conjugated fatty acids compared to WT breast tumors. This indicates that absence of CD47 in breast tumors is associated with a decrease in fatty acid availability. This was further validated in EMT-6 cells suggesting that blockade of CD47 reduces dependency on fatty acid oxidation compared to control cells.

The absence of CD47 resulted in a reduction of carcinogenesis, which was consistent with our previous studies and others examining antibody blockade of CD47 in a carcinogen-induced model ^51^. Because of our previous studies, we presume that the regulation of metabolism may occur autonomously in cancer cells; however, metabolites can influence inflammation in the tumor microenvironment resulting in the regulation of immunosuppression. Metabolites associated with tryptophan metabolism are linked to inflammation and immunosuppression in the tumor microenvironment. We observed a reduction of kynurenate, converted from tryptophan by the enzyme IDO, within *cd47*-/- breast tumors compared to WT tumors. Increased IDO activity is implicated in immunosuppression and resistance to immune checkpoint blockade ^52–55^. Our spatial proteomic analysis showed elevated immunosuppressive signature in WT breast tumors compared to *cd47*-/- breast tumors. We also observed the expression of the AhR receptor in WT mice. Since kynurenate binds to AhR, the reduction of both of these proteins suggests a potential mechanism for regulating immunosuppression regulated by CD47.

The most upregulated suppressive protein regulated in WT breast tumors compared to *cd47*-/- breast tumors were PD-L1. Both PD-L1 and CD47 expression have been previously implicated in resistance to cancer therapy, primarily due to the regulation of these genes by hypoxia ^55^. To determine if PD-L1 may interfere with the therapeutic efficacy of CD47 blockade, we combined these treatments in an orthotopic EMT-6 TNBC mouse model. Our data indicated that administering these therapies in combination caused a further reduction in tumor burden compared to anti-PD-L1 treatment or CD47 alone, suggesting a potential mechanism to enhance the efficacy of these drugs. Using these therapies in combination also caused a significant increase in infiltrating CD8+ T cells with improved antitumor function due to increased secretion of granzyme B compared to monotherapies and control, supporting previous studies ^12^.

The FDA has recently approved immune checkpoint blockade therapy, specifically targeting PD-L1, to treat metastatic TNBC. Thus, this data demonstrates how CD47 can be a potential biomarker and therapeutic target to enhance anti-PD-L1 therapeutic response for TNBC due to alterations in intratumoral proliferation, metabolism, inflammation, and immunosuppression. Clinical trials targeting CD47 have been conducted in patients with hematological and solid tumor cancers like non-Hodgkin’s Lymphoma, acute myeloid leukemia, and colon cancer ^5, 6^. Therefore, since these drugs are already in clinical studies, it would be feasible to include them in the invasive or advanced TNBC clinical space to reduce tumor burden and increase patient survival.

## Acknowledgments

D.R.S-P is supported by the NCI R21 (CA249349) P30 Pilot Breast Cancer Center of Excellence and the American Cancer Society Research Scholar Grant (133727-RSG- 19-150-01-LIB). E.R.S is supported by the NIGMS NRSA T32 Fellowship T32GM127261. S.M.B is supported by the NIH Office of the Director T32 in Comparative Medicine (T32 OD010957).

## Supplementary Materials

### Supplementary Figure Legends

**Supplementary Figure 1:**
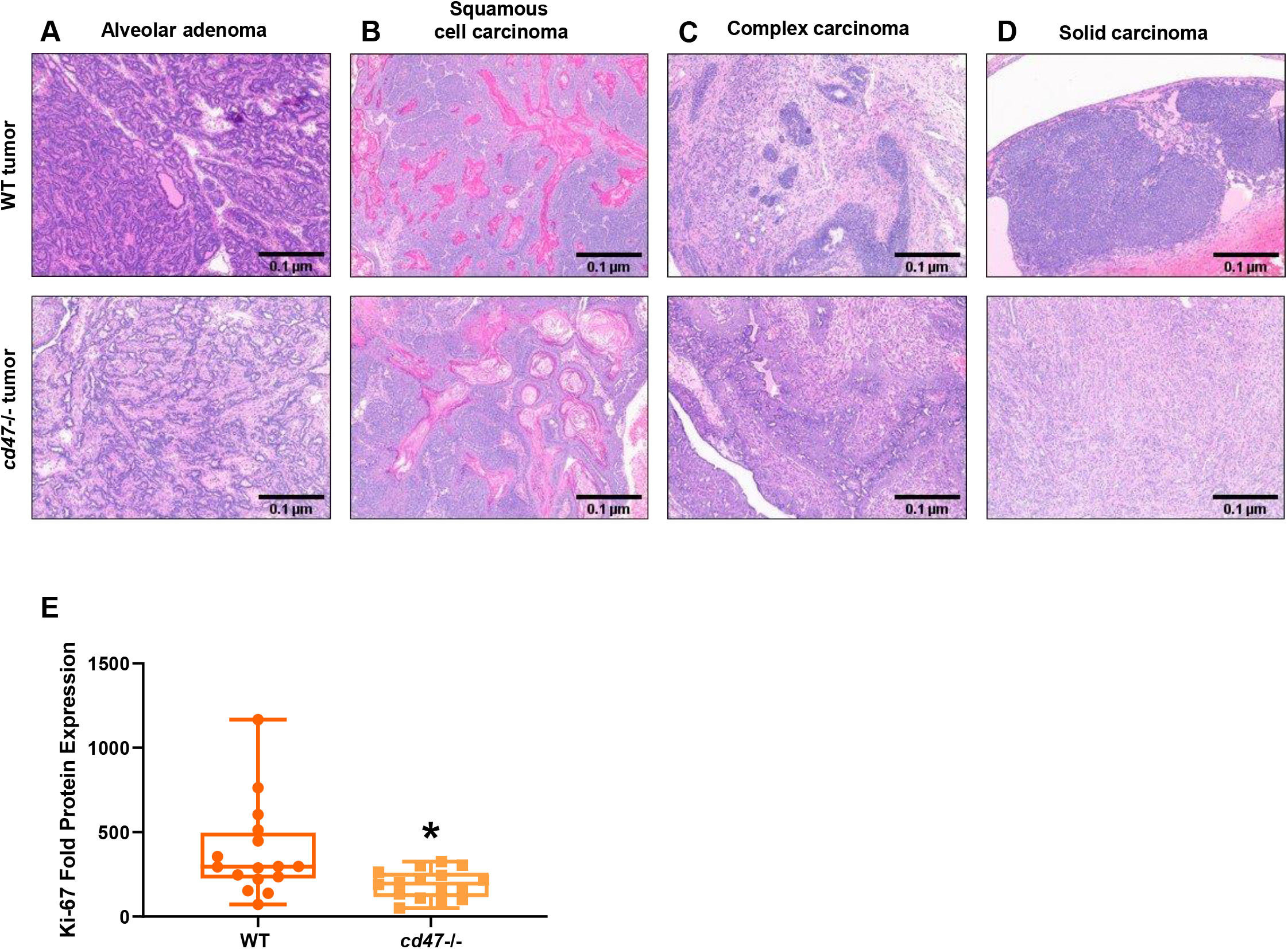
Hematoxylin and eosin staining of carcinogen-induced breast tumors were used to characterize tumors of WT and *cd47*-/- mice as **(A)** alveolar adenoma, **(B)** squamous cell carcinoma, **(C)** complex carcinoma, and **(D)** solid carcinoma. Images were obtained through light microscopy on the Olympus BX43 microscope at 10x magnification. Descriptions are based on the mouse mammary pathology consensus report ^56^. **(E)** Ki67protein expression decreased in *cd47*-/- breast tumors compared to WT breast tumors through spatial proteomic analysis (*p<0.05, 4/group).

**Supplementary Figure 2:**
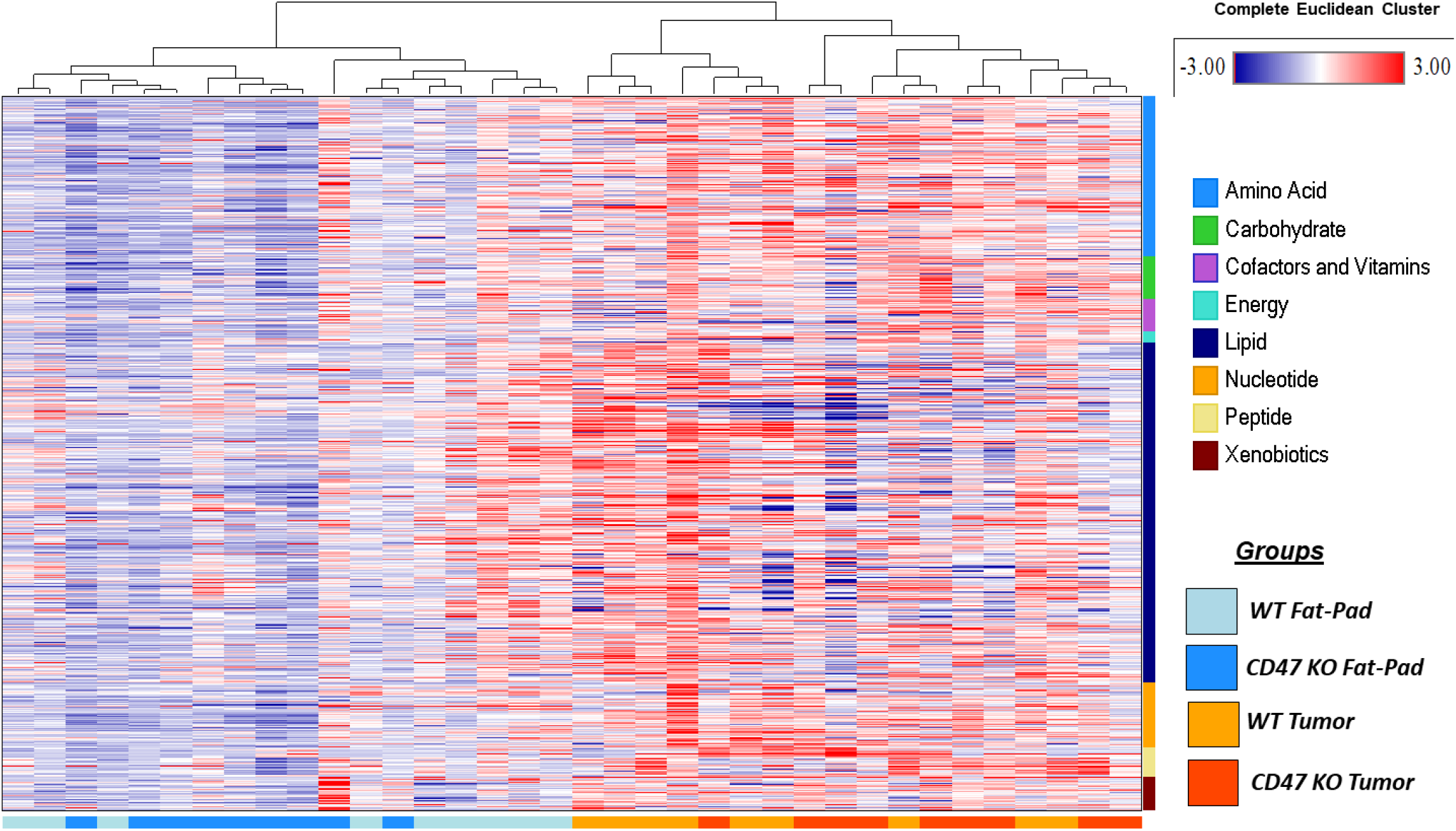
Hierarchical Clustering Analysis comparing metabolites in *cd47*-/- vs. WT mammary gland and tumors.

**Supplementary Figure 3:**
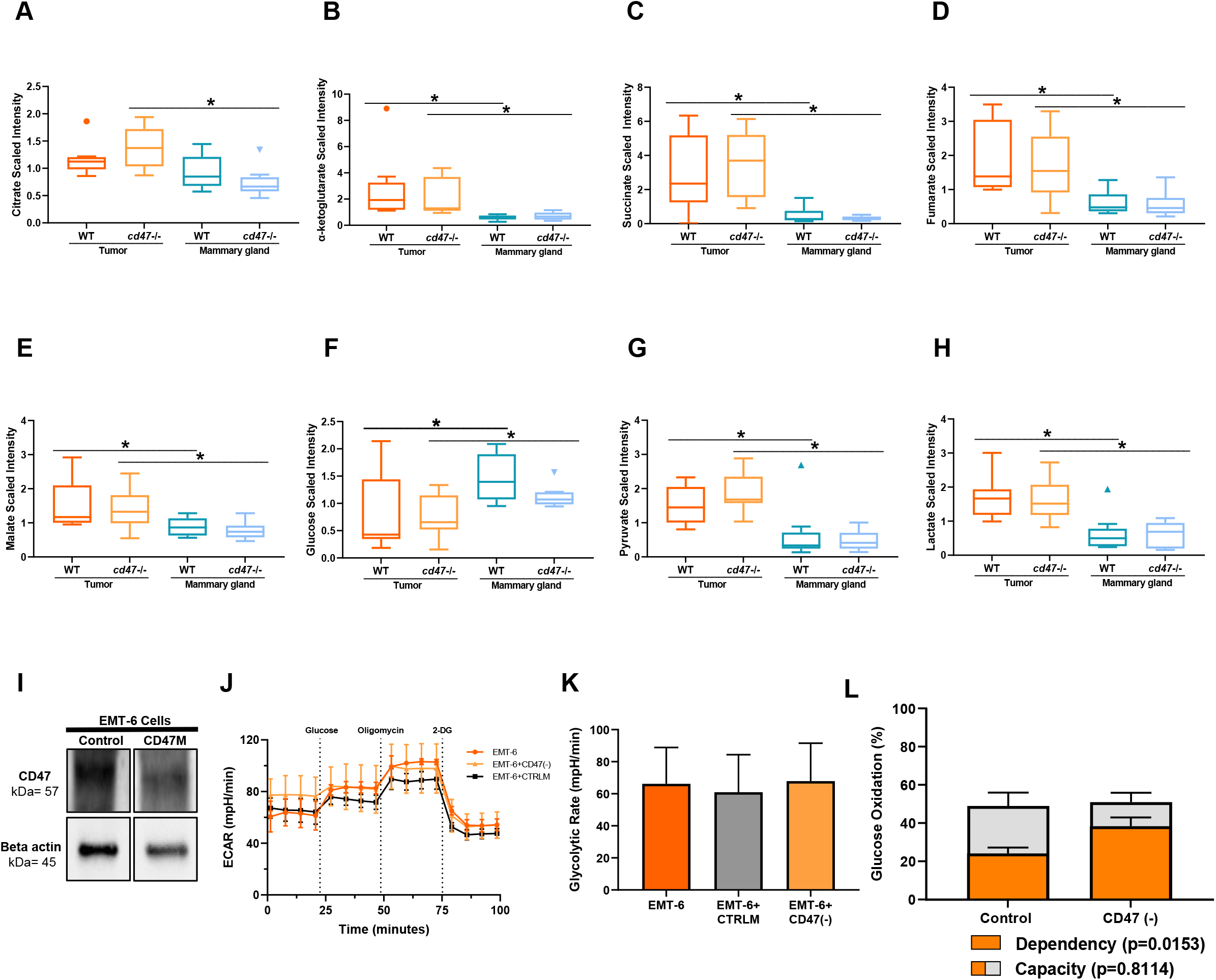
Breast tumors have an increase in metabolites associated with the citric acid cycle like **(A)** citrate, **(B)** α-ketoglutarate, **(C)** succinate, **(D)** fumarate, and **(E)** malate compared to mammary glands. A decrease in **(F)** glucose and an increase in **(G)** pyruvate and **(H)** lactate was observed in breast tumors compared to WT breast tumors. (*p<0.05, n=9/group) **(I)** CD47M decreases CD47 expression on EMT-6 cells. **(J)** Representative glycolytic flux of CD47 targeted EMT-6 cells. **(K)** No changes in the glycolytic rate but **(L)** an increase in glucose oxidation dependency in CD47 targeted EMT- 6 cells compared to control EMT-6 cells (*p<0.05, n=3).

**Supplementary Figure 4:**
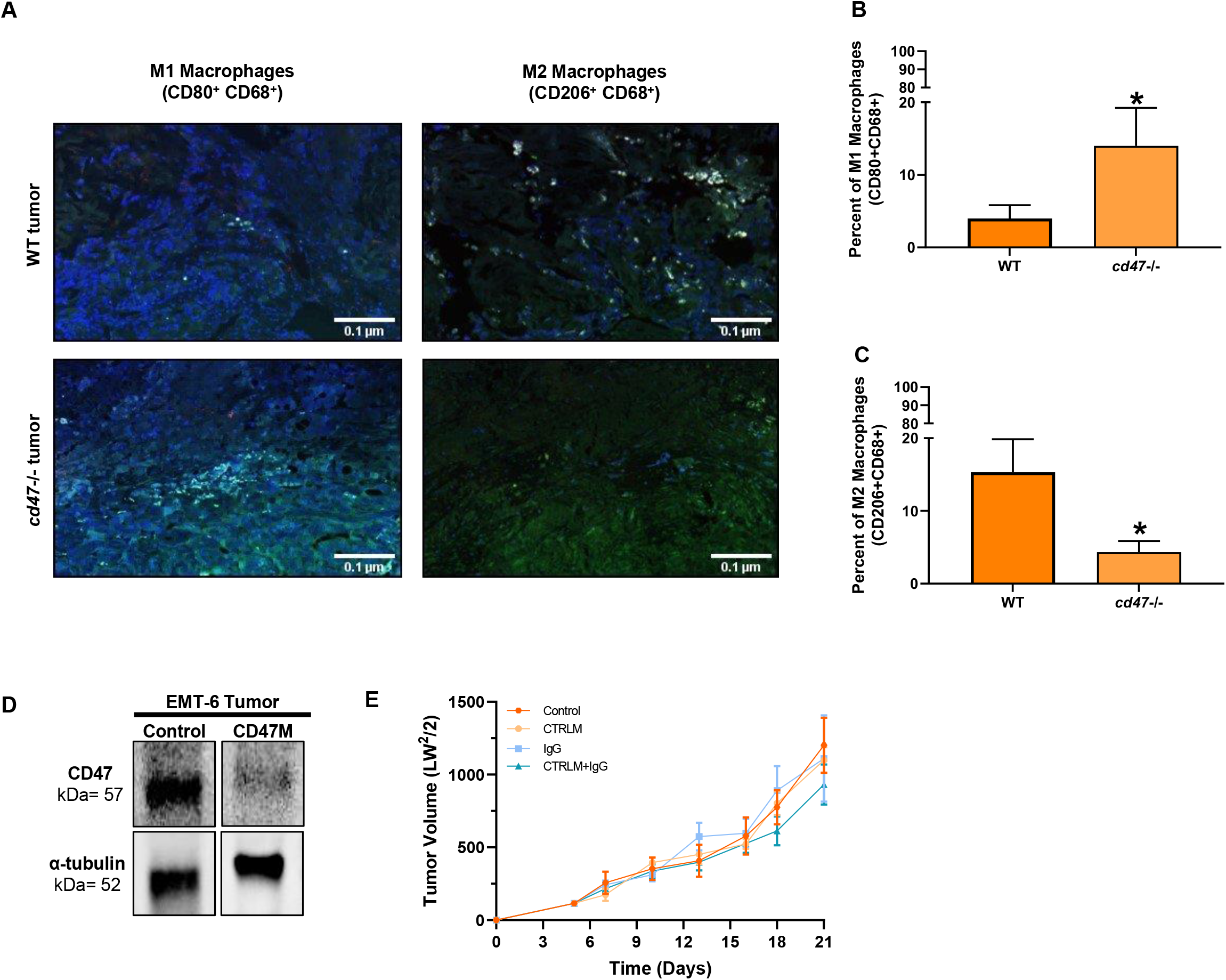
**(A)** Representative immunofluorescence staining to look at macrophage polarity within breast tumors. *cd47*-/- breast tumors have an increase in **(B)** M1 (CD80+ (PE red), CD68+(FITC, green), nuclei (DAPI, blue)) macrophages and decrease in **(C)** M2 (CD206+ (PE red), CD68+(FITC, green), nuclei (DAPI, blue)) macrophages compared to WT breast tumors (*p<0.05, n=3-4/group). **(D)** Decreased CD47 expression in EMT-6 tumors treated with CD47M. **(E)** There was no significant difference in the tumor volume of IgG and CTRLM treated mice compared to control (n=4- 7/group).

### Supplementary Table Legends

**Supplementary Table I:** Fold change of metabolites associated with the citric acid cycle.

|  |  | Two-way ANOVA Contrasts |  |  |  |
| --- | --- | --- | --- | --- | --- |
|  |  | Fold of Change |  |  |  |
|  |  | Tumor Mammary Gland |  | cd47 -/- WT |  |
|  |  | WT | cd47 -/- | Tumor | Mammary Gland |
| Citric Acid Cycle | Citrate | 1.26 | 1.84 | 1.17 | 0.80 |
|  | Aconitate [cis or trans] | 1.37 | 1.85 | 1.10 | 0.81 |
|  | Isocitrate | 1.88 | 2.92 | 1.43 | 0.92 |
|  | Alpha-ketoglutarate | 4.77 | 3.28 | 0.83 | 1.20 |
|  | Succinylcarnitine (C4-DC) | 0.61 | 0.60 | 1.06 | 1.08 |
|  | Succinate | 6.11 | 10.82 | 1.16 | 0.66 |
|  | Fumarate | 3.12 | 3.08 | 0.92 | 0.93 |
|  | Malate | 1.77 | 1.81 | 0.90 | 0.88 |
|  | Itaconate | 19.49 | 11.22 | 0.41 | 0.72 |
|  | Tricarballylate | 2.03 | 1.37 | 1.19 | 1.76 |
|  | 2-methylcitrate/homocitrate | 1.96 | 1.42 | 0.96 | 1.33 |

**Supplementary Table II:**
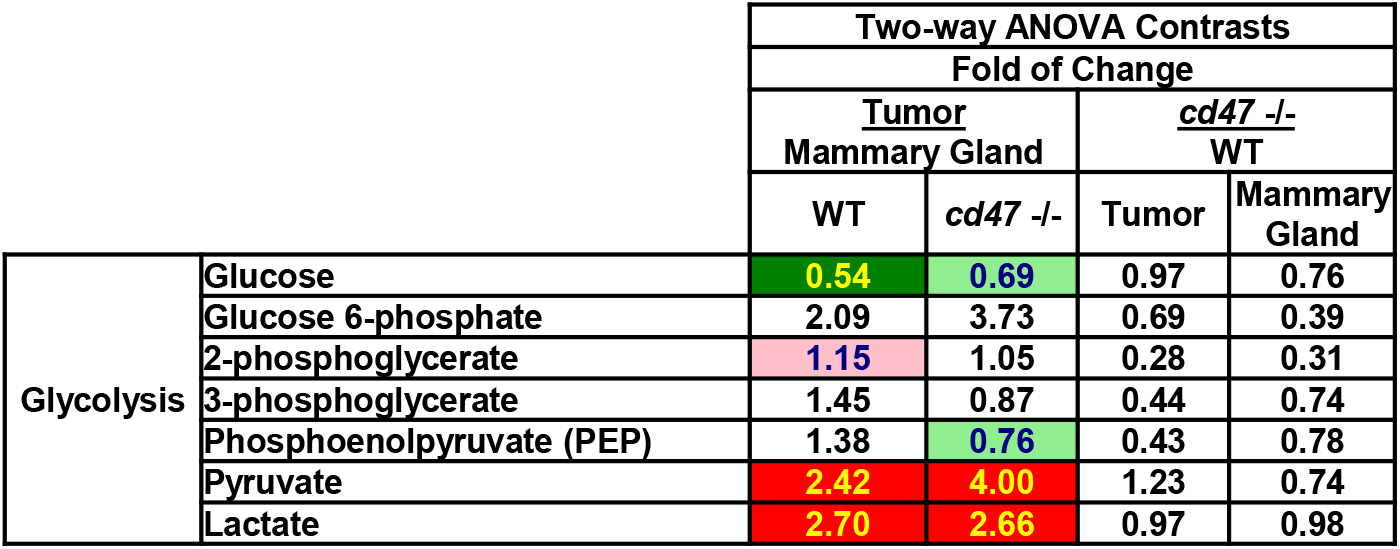
Fold change of metabolites associated with glycolysis.

**Supplementary Table III:** Fold change of metabolites associated with glutathione metabolism.

|  |  | Two- way ANOVA Contrasts |
| --- | --- | --- |
|  |  | Fold of Change |
|  |  | cd47 -/- WT |
|  |  | Tumor |
| Glutathione Metabolism | Glutathione, reduced (GSH) | 8.87 |
|  | Glutathione, oxidized (GSSG) | 2.71 |

